# Geometrical frustration of phase-separated domains in *Coscinodiscus* diatom frustules

**DOI:** 10.1101/2021.12.10.472144

**Authors:** Maria Feofilova, Silvan Schüepp, Roman Schmid, Florian Hacker, Hendrik T. Spanke, Nicolas Bain, Katharine E. Jensen, Eric R. Dufresne

## Abstract

Diatoms are single-celled organisms with a cell wall made of silica, called the frustule. Even though their elaborate patterns have fascinated scientists for years, little is known about the biological and physical mechanisms underlying their organization. In this work, we take a top-down approach and examine the micron-scale organization of diatoms from the *Coscinodiscus* family. We find two competing tendencies of organization, which appear to be controlled by distinct biological pathways. On one hand, micron-scale pores organize locally on a triangular lattice. On the other, lattice vectors tend to point globally toward a center of symmetry. This competition results in a frustrated triangular lattice, populated with geometrically necessary defects whose density increases near the center.

## INTRODUCTION

Phase separation is widely appreciated as an important mechanism used by cells to control biochemical pathways [1, 2]. However, cells also exploit phase separation to form functional structures with remarkable order and symmetry. For example, the keratinaceous color-producing nanostructures of bird feathers [3, 4], the topography of pollen’s cellulosic cell wall [5], and the porosity of silicaceous diatom frustules [6–8].

Diatoms are single-celled organisms with a beautifully structured cell wall made of silica, called the frustule. Their elaborate patterns have fascinated scientists and artists for hundreds of years [9]. The diatom builds its wall by precipitating silica in a specialized organelle called the silica deposition vesicle (SDV) [10] with the aid of biopolymers such as long-chain polyamines (LCPA) and silaffins [11]. While much progress has been made in understanding the chemical processes underlying silica deposition, little is known about the frustule’s organization at larger length scales. Experiments *in vitro* have shown that appropriate mixtures of LCPAs, silaffins, and other proteins can spontaneously form a porous silica structure [12, 13]. However, these structures are highly disorganized, and show none of the long-range organization that characterizes frustules. The organization of these structures has been hypothesized to either emerge spontaneously through the self-assembly of phase separated LCPA-rich domains [7] or through templating by the cytoskeleton [14, 15].

Here, we quantify the structural arrangement of micron-scale pores in valves of the diatom *Coscinodiscus granii*. We find that the pores locally form a triangular lattice. Globally, however, the structure is radially aligned. Other diatoms from the *Coscinodiscus* family also tend to form a local triangular lattice, but with varying degrees of radial alignment. Valves of *C. granii* grown in the presence of germanium maintain radial alignment in the absence of significant hexagonal order. Local hexagonal order is consistent with the self-assembly of mobile uniformly sized domains with steric repulsions. The mechanism underlying radial alignment is unknown, but it is apparently modulated across species. Using a simple Brownian dynamics simulation with hard discs and bond rotation, we created triangular lattices with various degrees of radial alignment. The simulated patterns span the range of morphologies observed in the *Coscinodiscus* family. While the competing tendencies of alignment and crystallization can readily be accommodated at the periphery of the frustule, geometric frustration leads to a highly disorganized core near the center.

## RESULTS AND DISCUSSION

### Pores of *C. Granii* make radially-aligned triangular lattice

We chose *Coscinodiscus granii* because its relatively large size and microscale pores enable optical microscopy for quantitative image analysis. In addition, it is one of the more established model diatoms [16–18]. To characterize the structure of valves, the flat regions on either end of the frustule, we imaged their micron-scale features using spinning-disk confocal microscopy (100x NA 1.45). Starting with 3-D image stacks of the extracted rhodamine-labeled diatom valves (Fig.1A), we calculated the 3-D centroids of the micron-scale pores in the valve (Fig.1B). From the side-view, it is clear that the valve is mostly flat, with some curvature near the edge. As locating pores was less reliable in these areas (shown in gray), we excluded them from further analysis.

For a given valve, the spacing between pores was very regular. The distribution of nearest-neighbor pore spacings from a representative valve is shown in Figure 1C. In this example, the average spacing obtained is 1.4 *μm*. While the overall size of valves varies by a factor of two (from 50 to 110 *μm* in diameter), the inter-pore spacing is very consistent, ranging from about 1.2 to 1.4 *μm* (Figure 1D). We observe a small but significant dependence of the pore-spacing with the size of the valve (Fig.1D,(slope 0.003, significant at *p <* 10^−8^). This suggests that pore size and spacing are largely determined by local processes (*e*.*g*. phase separation), independent of those that control the overall valve dimensions.

**FIG. 1:**
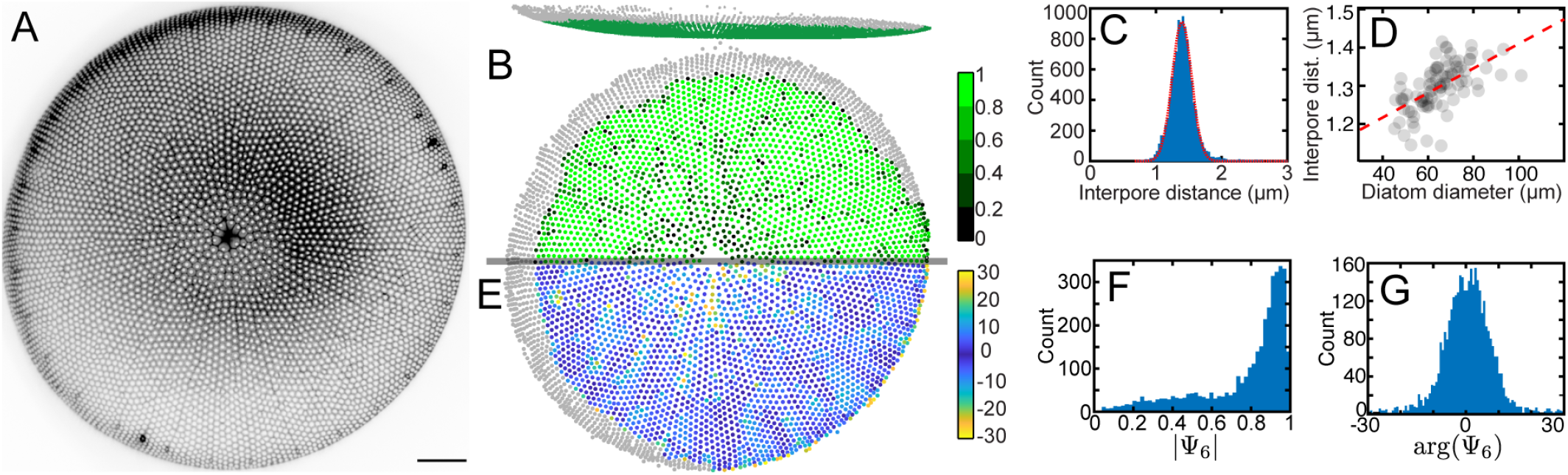
*Coscinodiscus granii* shows an evenly spaced pattern of micropores (A) An average intensity projection of a confocal image stack for a representative valve, scale bar 10 *μm*. (B) Reconstructed micropore structure. Top: Side view. Bottom: top view. Color represents absolute value of the radial Ψ_6_ parameter. Pores were located in 3-D using an automated procedure, areas of high slope (shown in grey) were filtered out. (C) Distribution of 3-D nearest-neighbor distance. (D) A weak dependence of pore-to-pore distance on diatom size is observed (slope 0.003 significant at *p <* 10^−8^). Each value is obtained by finding the peak of the distribution of inter-pore distance by fitting a gaussian curve. (E) Plot of the phase of the radial Ψ_6_ parameter. (F) Histogram of the absolute value of Ψ_6_. (G) Histogram of the phase of Ψ_6_.

Micropores show an intricate arrangement in the valve. In the vicinity of a single pore, the structure closely resembles a triangular lattice. However, a straight line drawn in any direction from the center to the edge of the valve will tend to follow the lattice.

To quantify this unusual structure, we adapted a geometrical tool developed for studies of 2-D crystals (see *e*.*g*. [19]). The hexatic order parameter, Ψ_6_, is defined with the complex expression

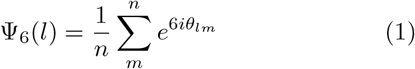

where *l* is a point on the lattice, and *m* is one of its *n* nearest neighbours (determined by Delaunay triangulation). *θ*_*lm*_ is the angle between the line connecting the *l*-th and *m*-th points, and a reference axis. The amplitude of Ψ_6_ is 1.0 for a perfect triangular lattice, but for a random arrangement of points we observe a distribution peaked at 0.3. Therefore, we can use |Ψ_6_| to locally assess crystalline order around each pore. This is visualized for one valve in Fig.1B, where each pore is colored according to |Ψ_6_|. A histogram of these values are shown in Fig. 1F. Most of the |Ψ_6_| values for this valve are close to one There is a very broad tail of low |Ψ_6_| values. These more disordered areas are more frequent near the center of the valve (Fig.1B).

On the other hand, the phase of the hexatic order parameter, arg(Ψ_6_), shows the local orientation of the lattice relative to a selected axis. Usually, a fixed axis is selected as a reference. Since our structure appears to be radially aligned, we found it more useful to select 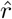, the unit vector connecting the center of the valve to the pore, as the reference axis for each point. By construction, this unusual choice of reference axis has no effect on |Ψ_6_|. However, it clearly shows how the local orientation of the crystal compares to 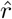. These orientations are shown point-wise for a typical valve in Fig. 1E. The distribution of arg(Ψ_6_), shown in Fig. 1G, is sharply peaked at zero. This demonstrates that the nearest neighbors of each micropore tend to be arranged in a hexagon that is oriented with one vertex pointing toward the center of the valve.

Thus, the pattern of holes in the valve can be thought of as the lattice constantly turning around the center of the valve. We define the overall crystallinity of a valve, 𝒞 = median(|Ψ_6_|). This will equal one for a perfect crystal. We define the overall radial alignment of a valve, 𝒜 = 1*/*variance(arg(Ψ_6_)). A well-aligned valve has narrow distribution of angles, and, therefore, a relatively large value of 𝒜.

A scatter plot showing the crystallinity and alignment parameters of *N* = 86 valves is shown in Fig.2A. Note that the values of 𝒜 and 𝒞 are correlated. This is counterintuitive, as crystallinity and radial alignment are incompatible. A closer look at the data shows that more highly organized valves tend to be larger, as indicated by the color-coding in Fig.2A.

### Lattice defects are polarized, inhomogeneous, and geometrically necessary

To further clarify the competition between alignment and crystallinity, we identified defect sites in the lattice. Using Voronoi tessellations, we determined the number of nearest-neighbors for each pore. In a triangular lattice, each point has six nearest neighbors. In a real 2-D crystal, there can be a significant number of defect sites with five or seven neighbors. The defect pattern of an exemplary valve is shown in Fig. 2C (defect patterns for additional valves are found in Fig. S8). There, points with six neighbours are shown in light gray, while five- and seven-fold defects are highlighted in red and blue, respectively. Most defects are found in five-seven (red-blue) defect pairs. These pairs are mostly isolated, only occasionally forming higher order structures. This is in contrast to 2-D crystals near equilibrium, where defect-pairs tend to condense into chains, forming boundaries between crystal grains.

**FIG. 2:**
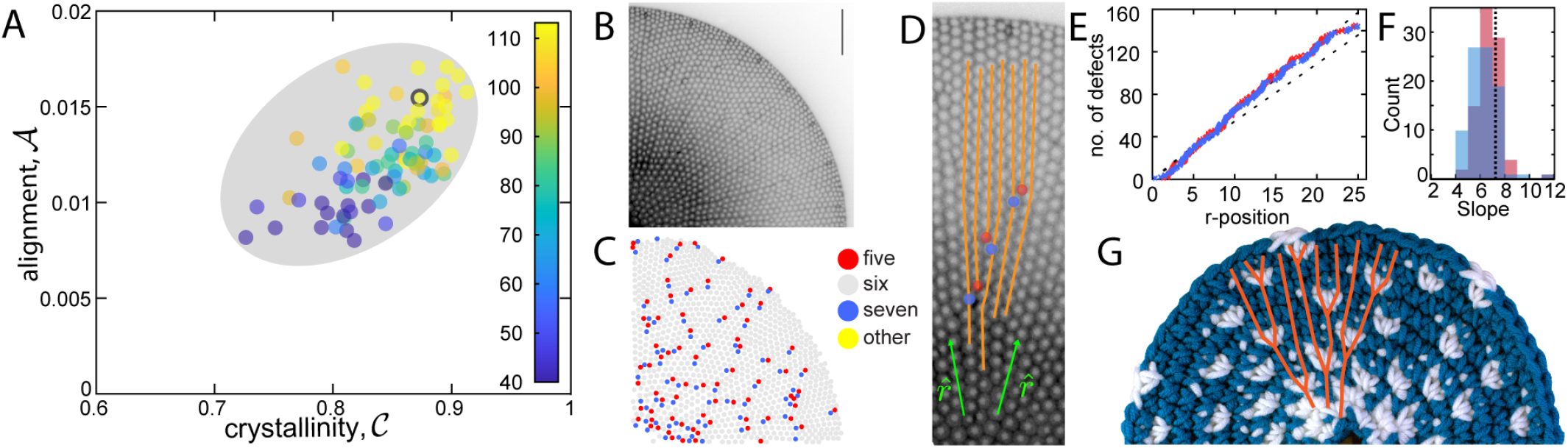
*C. granii* diatoms display different alignment, crystallinity and defect patterns. (A) Radial Ψ_6_ phase-amplitude plots, with gray oval to guide the eye. Datapoint color shows diatom size in *μm*. Larger diatoms tend to be both more aligned and more crystalline. (B) Example of a confocal stack projection of the diatom labeled in A. Scale bar 10*μm*. (C) Tracked pores of the same diatom, showing points with 6 neighbours in gray, 5 in red and 7 in blue. Pairs of 5-7 (red-blue) defects are scattered over the structure, with 5 (red) mostly located closer to the edge and 7 (blue) towards the center. (D) Radial lined traced over a section of a confocal frustule image. Defects shown in red (5) and blue(7). Radial direction indicated in green. (E) Cumulative number of defects (7-connected shown in blue, 5-connected in red) plotted against their radial position. Dashed line shows slope 7.2. (F) A histogram of slopes for all observed diatoms, same color scheme. Vertical dashed line is showing 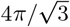 (G) Photograph of a crochet circle, with the increase locations shown in white.

Intriguingly, defect pairs are polarized, with the seven-fold defect closer to the center than the five-fold one. Note that five-seven defect pairs correspond to the termination of a crystal line, known as an edge dislocation, illustrated in Fig. 2D [20]. To avoid overlapping pores, crystal lines must terminate as they converge on the center of symmetry.The five-fold defect sits at an end of the terminated crystal line, and the seven-fold defect is found just beyond it. The observed polarization of the defect pairs is therefore a direct consequence of radial alignment of the lattice vectors.

Defects are more frequent near the center of the valve. To quantify this, we plot the cumulative number of defects within a distance, *r*, from the center for an exemplary valve in Fig. 2E. For a conventional 2-D crystal, the defects are distributed homogeneously, and their cumulative number would increase as *r*^2^. However, for diatom valves, the cumulative number increases with linearly with *r*. The mean slope is 6.8 *±* 0.1 and 6.3 *±* 0.1 defects per lattice spacing for five- and seven-fold defects, respectively (Fig. 2F). Note that a linear scaling of the cumulative number of defects is indicative of a density scaling like 1*/r*.

While radially-aligned crystal structures are unusual in crystalline materials, they are common in the traditional craft of crochet, as shown in Fig.2G. In a circular crochet, the square crochet lattice grows radially, because each new stitch is inserted, by default, in a stitch on the previous row. However, every crocheter knows that each new row must be expanded by approximately six stitches for the circle to lay flat [21]. This is geometrically necessary, as the circumference of each layer increases by 2*πa* for each row, where *a* is the spacing of the stitches.

The same argument applies to the diatom lattice, but the increase in number of lattice sites per row is slightly different because the underlying lattice is triangular (see Supplement). When the triangular lattice is aligned with the radial direction, the defect insertion rate is 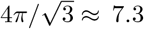 defects per lattice spacing. This value is comparable to the range of values we observe, as shown in Fig. 2F.

### Crystallinity and Alignment are Controlled by Distinct Biological Mechanisms

So far, we’ve focused on structural analysis of a single species, *C. granii*. The morphology of diatom frustules, however, is tremendously diverse. Within the *Coscinodiscus* family, frustules are generally barrel-shaped with flat round valves featuring organized patterns of microscopic pores. We collected two-dimensional images of the valves of other *Coscinodiscus* diatoms from a variety of sources, shown in Fig. 3A. We located pore positions using a semi-automated procedure (Fig.3B), and analysed them the same way as our *C. Granii* data. The crystallinity and alignment parameters for these diatoms are shown in Fig. 3C. While samples from *C. radiatus* and *C. wailesii* fall into the range of morphological parameters we observed in *C. granii*, samples from *C. angustelineatus* and *C. eccentricus* are completely different. They show excellent hexagonal order, with no indication of radial alignment.

**FIG. 3:**
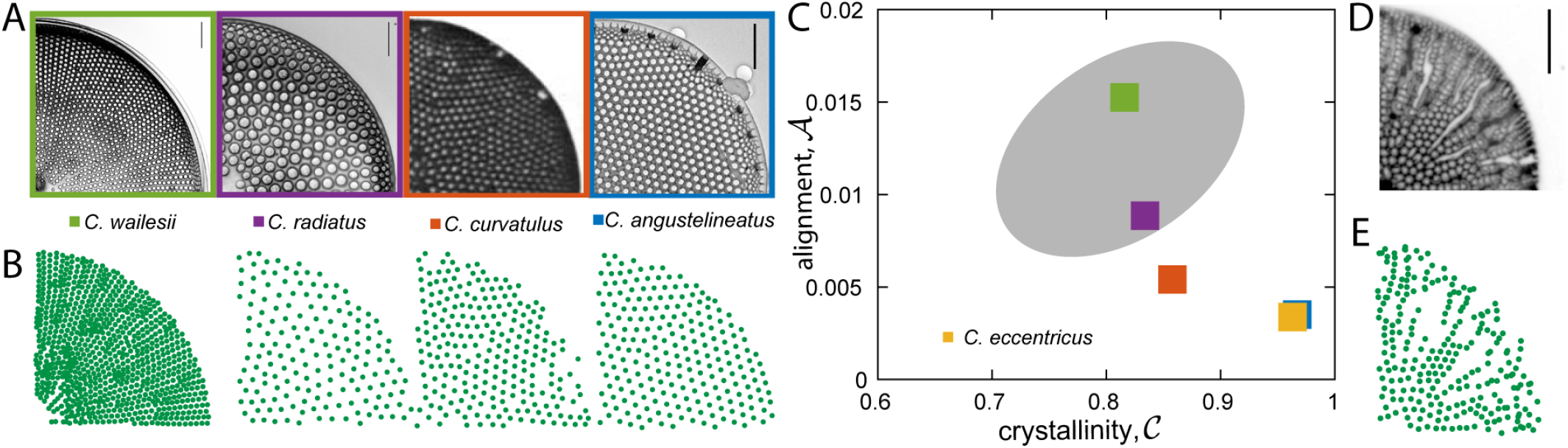
*Coscinodiscus* family diatoms. (A) Images of diatoms. *C. angustelineatus* and *C. eccentricus* (see S3) from [22] (Reprinted by permission of Taylor & Francis Ltd). *C. curvatulus* from [23] (©The Royal Society of New Zealand, reprinted by permission of Informa UK.), *C. wailesii* and *C. radiatus* provided by Dr. Johannes Goessling. Scale bar 10*μm*. (B) Corresponding points obtained using the locating procedure. (C) Radial Ψ_6_ phase-amplitude plots of coscinodiscus family diatoms shown in A, with gray oval representing *C. granii*. (D) Confocal stack average projections of *C*.*granii* diatoms grown in presence of germanium. Scale bar 10*μm*. (E) Corresponding points obtained using the locating procedure.

For *C. granii*, alignment of pores is very robust, even when crystallinity is destroyed by elemental substitution. The structure of the silica frustule can be altered by growing diatoms in the presence of germanium [24, 25]. Since germanium and silicon belong to the same group of the periodic table, they have similar chemistries. When germanium is introduced into the cell-culture medium, it is taken up by the cell and incorporated into the frustule. Over a range of germanium concentrations, diatoms continue to grow and divide at a normal rates (Fig. S4). However, the morphology of the frustule can change dramatically, as shown in Fig. 3D and S5. Elemental imaging (LA-ICP-TOFMS) indicated a distribution of germanium at a uniform rate across the frustule (Fig. S6). While the morphology of germanium-perturbed valves changes significantly from cell-to-cell, the pores tend to have a wider range of sizes and shapes, which include crack-like gaps in the frustule. In these conditions, crystalline order has essentially disappeared. Nevertheless, strong alignment of pores along the radial direction is evident in the micrographs (*e*.*g*. Fig. 3D).

We have found that the organization of pores in diverse natural and perturbed valves are characterized by a blend of crystallinity and radial alignment. On one hand, natural variations across the *Coscinodiscus* family maintain crystallinity, but feature varying degrees of radial alignment. On the other, perturbation of *C. granii* with germanium appears to maintain radial alignment while destroying crystallinity. This suggests that crystallinity and alignment are driven by different mechanisms.

### A Colloidal Model with Bond-rotation Captures the Diversity of Observed *Coscinodiscus* Structures

Since the biochemical mechanisms that regulate these patterns are unknown, we developed a minimal physical model to understand the competition of radial alignment and crystallinity. Crystallinity emerges naturally in two-dimensional systems of Brownian particles with repulsive interactions. We consider a set of *N* disks with radius *a*, placed randomly within a circular confining potential. The particles diffuse, and repel each other with a force proportional to their overlap, quantified with a spring constant *k*_*r*_. In addition, we introduce an alignment force that tends to rotate the bond between neighboring particles to align with the local radial direction. The magnitude of the alignment force also depends on the degree of overlap, and is proportional to a spring-like constant, *k*_*a*_. For more details, see the methods section.

We performed a series of Brownian dynamics simulations with a varying strength of alignment, *k*_*a*_, and fixed repulsion, *k*_*r*_. Particles were randomly distributed within the circular simulation domain with an effective area fraction *ϕ* = *N* (*a/R*)^2^ = 1.1, and evolved in time until reaching steady state. Two system-sizes were considered, with *N* = 1760 and *N* = 7948, roughly matching the smallest and largest number of pores observed the valves of *C. granii*.. The resulting steady structures are shown in Fig. 4A-C. The crystallinity and alignment parameters of the resulting structures are shown in Fig. 4D. As *k*_*a*_ increases, the crystallinity parameter drops and the alignment parameter increases. While alignment increases only modestly for the smaller system size (triangles), it increases much more strongly for the larger system size (circles). At large values of *k*_*a*_, the simulated systems achieve values of crystallinity and alignment typical of *C. granii* valves. For a fixed value of *k*_*a*_, the larger valves show better crystallinity and alignment parameters, echoing the size-dependence of real valves, shown in Fig. 2A. Fig. 4E shows the cumulative count of defects as a function of distance from the center for *k*_*a*_ = 0 and 0.8. Like in real diatoms, the simulated structures at *k*_*a*_ = 0.8 have a linearly decreasing defect density with distance from the center, with a slope approaching 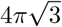, indicated by the dashed line. On the other, defect-pairs in the simulation tend to form higher-order structures defining more pronounced grain-boundaries than observed in the real *C. granii* valves, suggesting that the latter structures may have kinetically arrested before finding their energetically preferred configuration [5].

**FIG. 4:**
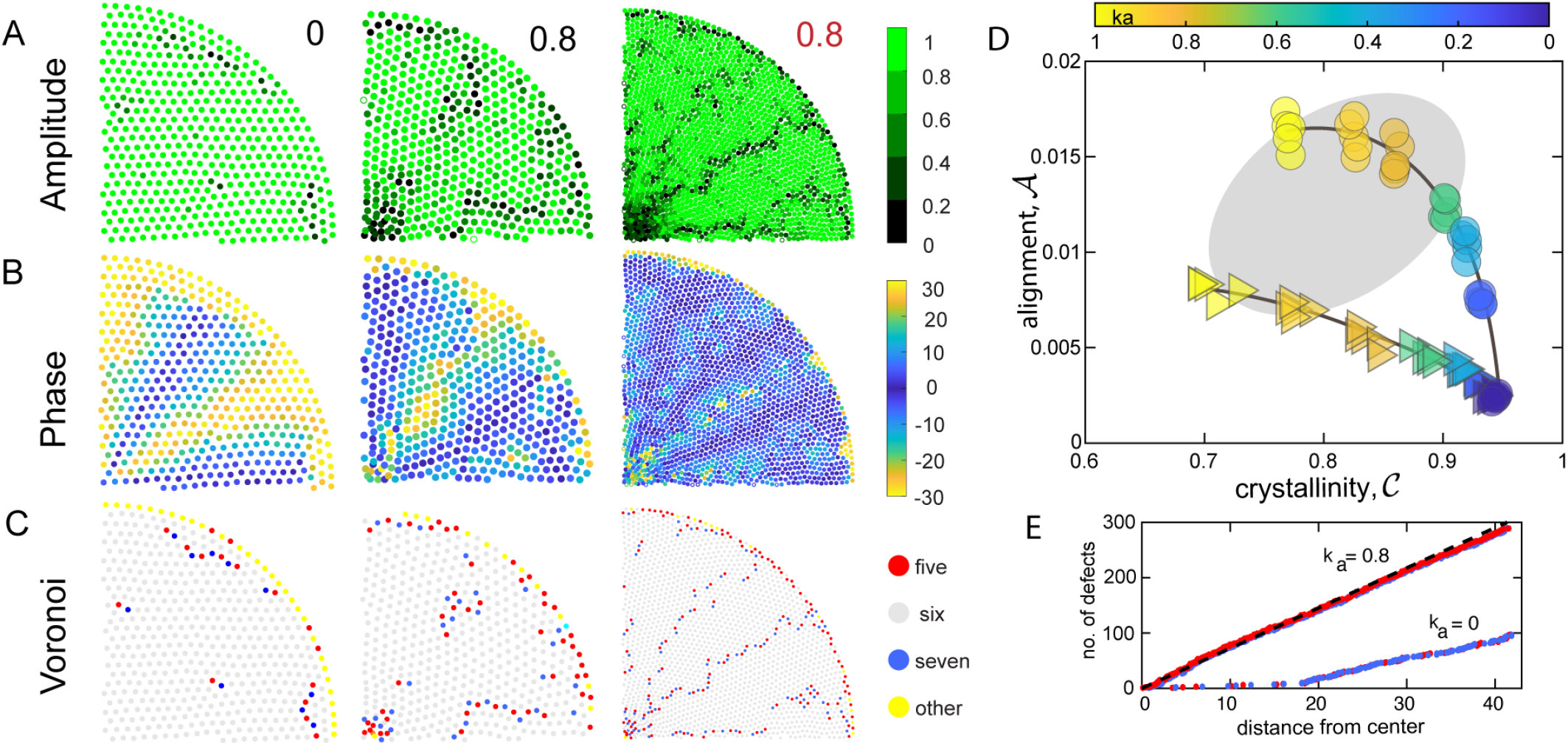
Simulations yield diatom-like structures. (A) Radial Ψ_6_ phase-amplitude plots, with gray oval representing *C. granii* data. Triangle color represents the value of *k*_*a*_ used in the simulation. For each value the simulation was run five times, for each run the initial distribution of points was randomized. (B) Examples of Ψ_6_ amplitude plots for several values of *k*_*a*_ for small and large numbers of particles. (C) Radial Ψ_6_ for the same data. (D) Color-coded representation of the number of neighbours. Points with 5 nearest neighbours are shown in red, 6 in gray, 7 in blue and others in yellow. (E) Cumulative number of defects plotted against distance from center for *k*_*a*_ = 0.8 and 0 for *N* = 7948. Five-fold defects plotted in red, seven-fold in blue. Dashed line shows slope 7.2.

## CONCLUSION

We have found that micropores of *Coscinodiscus granii* diatom valves are arranged on radially-aligned triangular lattices. These novel structures feature a geometrically necessary increase of defect density near the center of the valve. While triangular lattices are observed across the *Coscinodiscus* family, the degree of radial alignment varies from species to species. When the crystallinity of *C. granii* is perturbed by the introduction of germanium, the valve structure remains highly aligned. These observations suggest that these two structural features have independent physical origins. The range of structures observed across the *Coscinodiscus* family can be recapitulated in simulations that include conventional steric repulsions as well as a torque aligning their separation vectors with the local radial direction.

While steric repulsions are consistent with the self-assembly of stable phase-separated domains, the origin of the torques producing alignment is unknown. At a minimum, this requires the imposition of a center of symmetry, which may be acheived by the actin cytoskeleton, whose filaments have shown to been radially aligned in close association with the outer surface of the SDV [14]. Alternatively, confinement of the SDV within the curved boundary of the cell could lead to radially aligned creases [26, 27]. Direct live-cell observations of frustule development will be essential to identify the mechanisms driving alignment, and more generally, to understand the phase separation process that produces micropores.

Whatever its origin, radial alignment imposes geometrically necessary defects in a flat two-dimensional lattice. Specifically, pairs of five- and seven-fold defects need to be inserted with a frequency inversely proportional to the distance from the center of symmetry. This is in stark contrast with two-dimensional crystals conforming to a curved surface in the absence of a center of symmetry. There, five- and seven-fold defects are introduced in an imbalanced manner, with a topological charge density proportional to the local Gaussian curvature [28, 29]. Traditional knowledge of crochet demonstrates that radial alignment in the absence of defects imposes curvature. Indeed, a defect-free radially symmetric crochet lattice deforms to produce a tube. Furthermore, an appropriate insertion of defects produces fascinating structures including hyperbolic planes [30] and Lorentz surfaces [31]. More generally, insertion of defects could provide a route to control the embedding of a 2-D object in 3-D space by specifying the local curvature [32–34].

Diatoms are master-builders of inorganic structures at the microscale. We expect that a deeper understanding of the processes they use to sculpt biomineralization could inspire new approaches to sustainable micro- and nano-manufacturing.

## Methods

### *Coscinodiscus granii* cells

*Coscinodiscus granii* (strain K-1834) diatoms were purchased from The Norwegian Culture Collection of Algae, Oslo, Norway and cultured in L1 media [35]. The cells were cultured in filter-top flasks in a temperature and light-controlled incubator (Tritech Research, Inc., Los Angeles, USA) at 15°C under a red and blue LED grow-light. The light was cycled 14h/8h on/off, light flux when on was 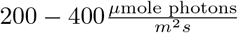.

Fluorescent cell walls were obtained by metabolic insertion of Rhodamine B (Sigma-Aldrich, St. Louis, USA). The Rhodamine dye was added to the culture medium at a concentration of 1*μM*, the culture then was grown for at least 24 hours [36]. Diatom cell walls were extracted using a “soft-cleaning” method based on [36]. Cells were collected by centrifugation at 2400*g* and re-suspended in 0.1*M* Na–EDTA and 2% sodium dodecyl-sulphate solution and heated at 90^*°*^*C* for 5 minutes.

### Imaging

Imaging was performed with a Nikon eclips-Ti microscope (Nikon Corporation, Tokyo, Japan), equipped with a spinning disk confocal attachment (Intelligent Imaging Innovations, Inc., Denver, USA). Imaging was performed with a Nikon PLANAPO 100x NA 1.45 lambda oil immersion objective or Nikon PLANAPO 100× NA 1.45 lambda oil immersion objective (Nikon Corporation, Tokyo, Japan) for germanium diatoms.

### Image analysis

Image analysis of *Coscinodiscus Granii* diatoms has been performed in MATLAB (MathWorks, Natick, MA), using an automated procedure. The image z-stacks were contrast adjusted, inverted, pre-processed with a bandpass filter and binarized using the MATLAB function `imbinarize`. For each object, area, and solidity was determined using `regionprops`, both were used for filtering. Z-positions were determined by first finding the track of each object through the z-stack following the method in [37] implemented for MATLAB in [38]. Then, for each object, the z-plane with the largest area was selected as the final z-coordinate. The center of the structure was determined manually.

Image analysis of other *Coscinodiscus species* was performed on 2D images using MATLAB. The images were pre-processed by being inverted (when necessary), filtered with a bandpass filter, thresholded and binarized. The function `regionprops` was used to locate features and remove noise by thresholding feature size. The center of the structure was determined manually.

### Elemental analysis

LA-ICP-TOFMS element imaging [39] was performed using the single pulse mode according to Neff et al. [40]. A laser ablation system (GeoLas C, Lambda Physik, Germany) employing a modified tube cell design [41, 42] were coupled to an ICP-TOFMS instrument (icpTOF2R, Tofwerk AG, Switzerland). The piezo-electrically driven xyz translational stage (1 nm scan resolution, SmarAct GmbH, Germany), the laser and the TOFMS data acquisition were triggered using a custom-built imaging control system[40]. The full mass spectrum from Na to U was acquired and the quantification of element mass fractions were performed for each pixel based on 100% mass normalization[43] using NIST SRM 610 as an external reference material and assuming that all elements are present as oxides.

### Simulations

Simulations were performed in MATLAB (Math-Works, Natick, MA).

Consider a set of *N* discs with radius *a*, located at 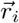, where *i* = 1, …, *N*. The equation of motion of a single object is given by

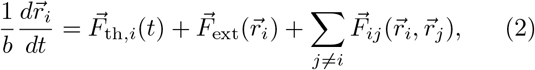

where *b* is the hydrodynamic mobility. Particles interact with a thermal bath, which generates a random force, 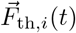. This force is uncorrelated in space and time, and drives Brownian motion. Particles are subject to a position-dependent external force, 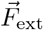. Here, we construct an external force to limit particle motion to a circular area of radius *R*. The center of this circular domain is taken to be the origin for all of the particle locations. Finally, particles interact with each other: 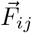 is the force exerted on the *i*^th^ particle due to the *j*^th^ particle.

Overlapping particles interact in two ways,

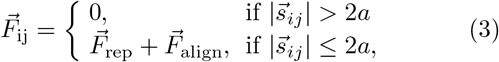

where, 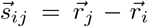 is the vector separating particles *i* and *j*. The first contribution to the interaction is a conventional spring-like repulsion,

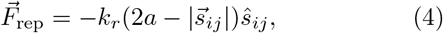

whose strength is determined by a spring constant, *k*_*r*_. With only this interaction term, sufficiently concentrated particles will evolve into a crystalline steady state with a triangular lattice. Therefore, we introduced a second interaction that works to point the separation vector toward or away from the center of the structure,

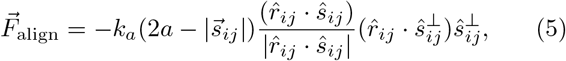

Where 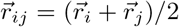. This force is applied perpendicular to the separation vector along 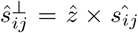, where 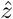 is the unit vector pointing out of the plane. Like the repulsive force, it depends on a spring constant, *k*_*a*_, and is proportional to the extent of overlap between the particles. However, instead of pushing the particles away from each other, it rotates them around their center of mass, 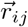. The strength of 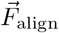 increases as 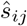 turns away from 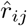. This specific form is not motivated by a specific physical model, but is simply meant to capture a generic tendency for alignment.

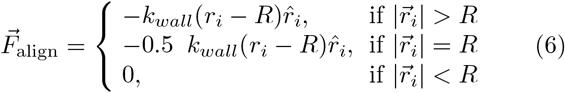

where *r*_*i*_ is the distance form the center of the structure to particle *i* and 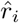 is the radial unit vector pointing from the particle to the center. The spring constant of the wall was 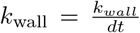 where *k*_*wall*_ = 1. The radius of the confining wall was *R* = 20 or 40 for *N* = 1760 or *N* = 7948 particles respectively. The filling fraction was kept constant at *ϕ* = 1.1. We give the particles a diffusion coefficient *D* = 0.01.

Simulations were run with a time-step of *dt* = 0.05 for a total number of time-steps *N*_*t*_ = 80000. The parameter *k*_*align*_ was varied from 0 to 1.2. The simulations were run five times for each value of *k*_*align*_, with initial particle positions randomized for each run.

## Acknowledgments

The authors would like to thank Prof. Detlef Günther and Dr. Christoph Neff for the elemental analysis data; Kapitalina Kuzmina for tutelage in yarn craft; Dr. Asja Radja, Dr. Robert Style, and Prof. Roel Dullens for great discussions; Dr. Johannes Goesseling for *Coscinodiscus* images and vital advice on setting up the diatom culture; Prof. Joseph Paulsen for pointing us to the thin sheet wrinkling hypothesis.

## Supplemental materials Defects in radial lattices

Simple crocheting produces a square lattice, because every new loop is built on a loop in the previous row. To tile a circle radially with stitches, double stitches, which form additional radial columns, must be introduced (Fig. 2G). It is possible to calculate the number of increases from geometrical considerations. The circumference of a row at radius *r* is *C* = 2*πr*. The difference in length between each new row and the previous row is

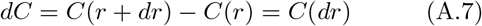

Let *h* be the height and *w* the width of each stitch (each tiling unit), then *dr* = *h* and the length difference in units of rows is *C*(*dr*) = 2*πh*. The difference in units of lateral stitches is 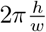. If the stitch is “square” (height is equal to width as shown in S1A), then 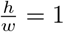 and

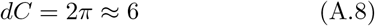

Thus, each row should have about 6 more stitches than the previous.

In the case of the diatom, the circle is tiled with a triangular lattice, and, according to our data, the triangles are turned such, that the edges are facing radially, as shown in Fig.S1C. In this case *h* = *a*, where *a* is the lattice spacing. The lattice consists of equilateral triangles, *w* is in this case the height of the triangle (Fig.S1D) and 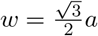. It follows,

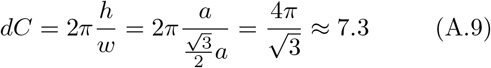

Each row must therefore have ≈ 7 increases. In the case of a triangular lattice, each increase creates a new radial line with a 5-7 defect pair at the start of the line, as shown in Fig.2D.

**FIG. S1:**
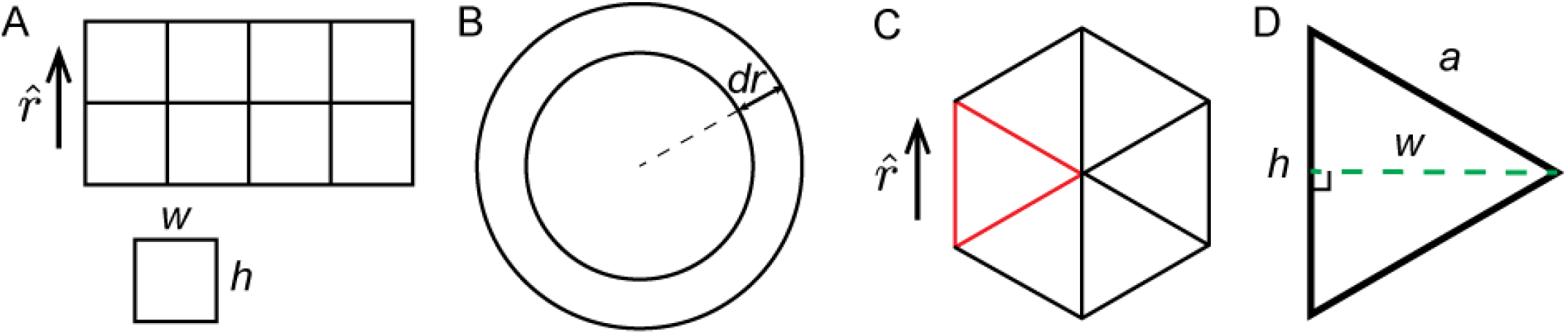
A sketch of the orientation of the triangular lattice.

**FIG. S2:**
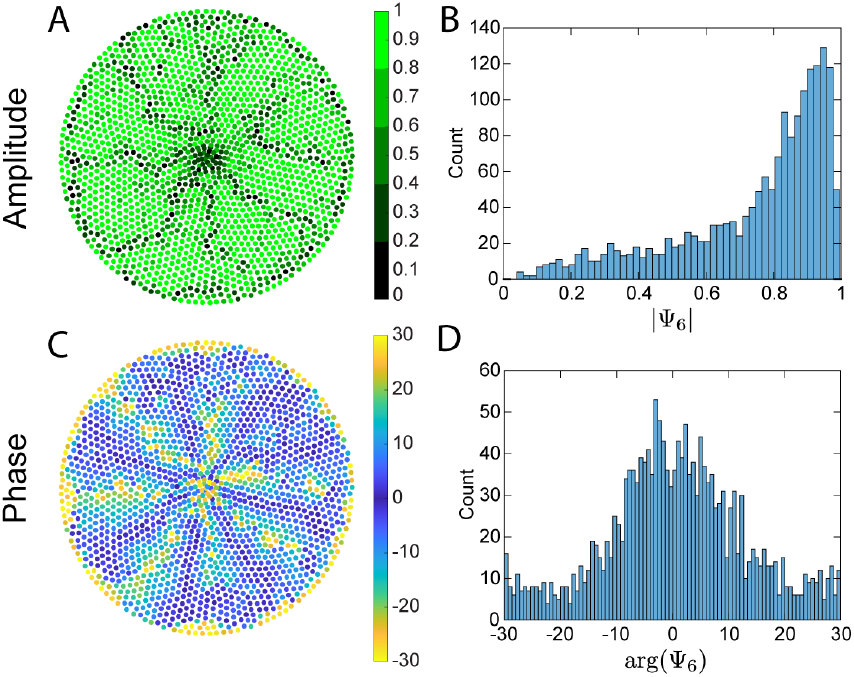
Ψ_6_ evaluation for simulation results. (A,C) amplitude and phase plots for *k*_*align*_ = 0.8. (B,D) respective histograms.

**FIG. S3:**
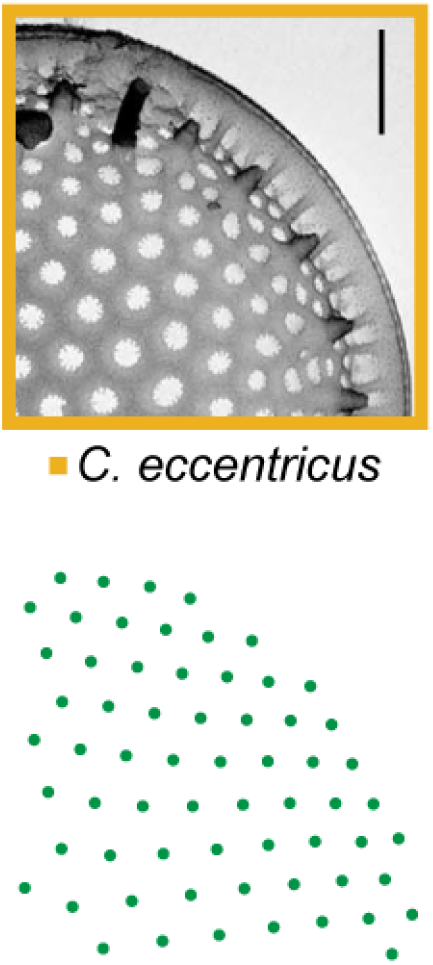
*C. eccentricus* image and tracked micropores. Image from [22] (Reprinted by permission of Taylor & Francis Ltd)

**FIG. S4:**
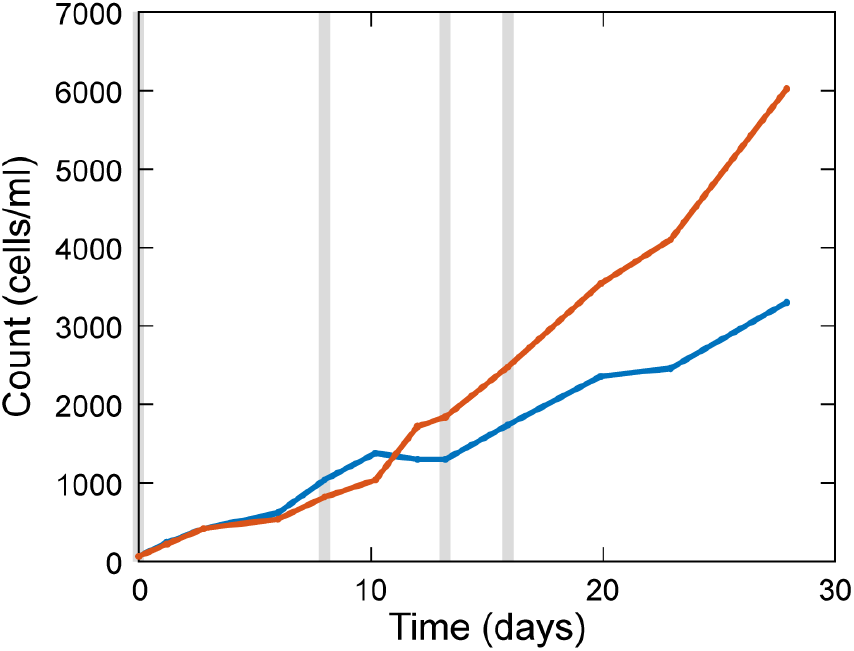
*C. granii* growth curves shown for germanium culture (red) and for control without germanium (blue). Gray vertical lines signify addition of feeding solution with 17 *μM* of Si in both solutions, the feeding solution with germaniun contained also 0.32 *μM* Ge (molar ratio Si:Ge = 53).

**FIG. S5:**
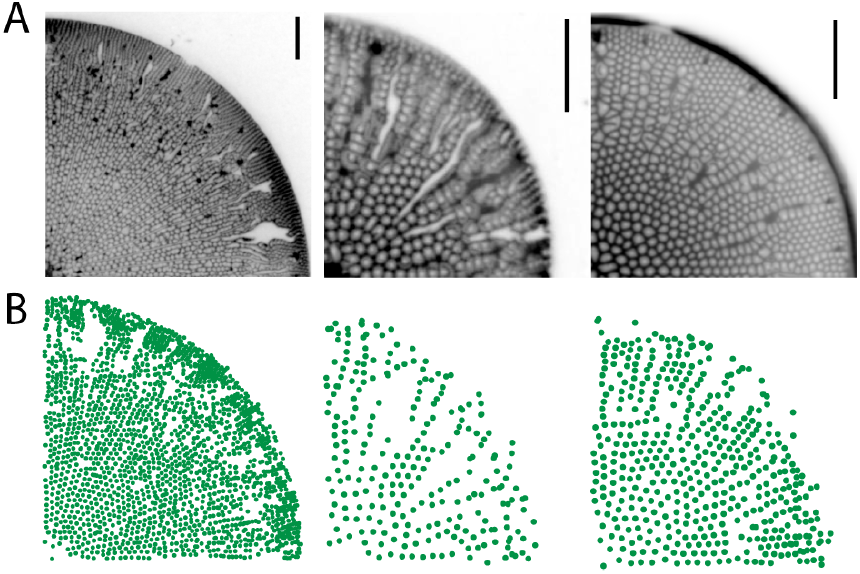
Ge

**FIG. S6:**
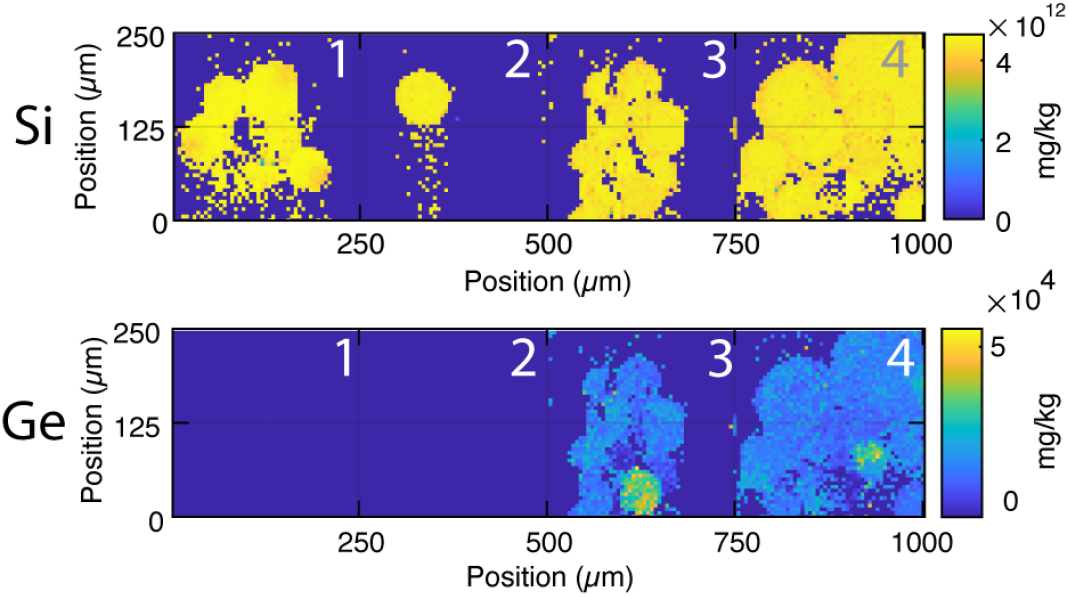
Element mapping of diatom frustules. Extracted frustules of: 1,2 *C. granii* cells grown in L1 media, 3, 4 *C. granii* cells grown in L1+germanium. Top row is Si detection, bottom row is Ge detection. The spatial distribution maps suggest, that germanium is distributed evenly over the frustule.

**FIG. S7:**
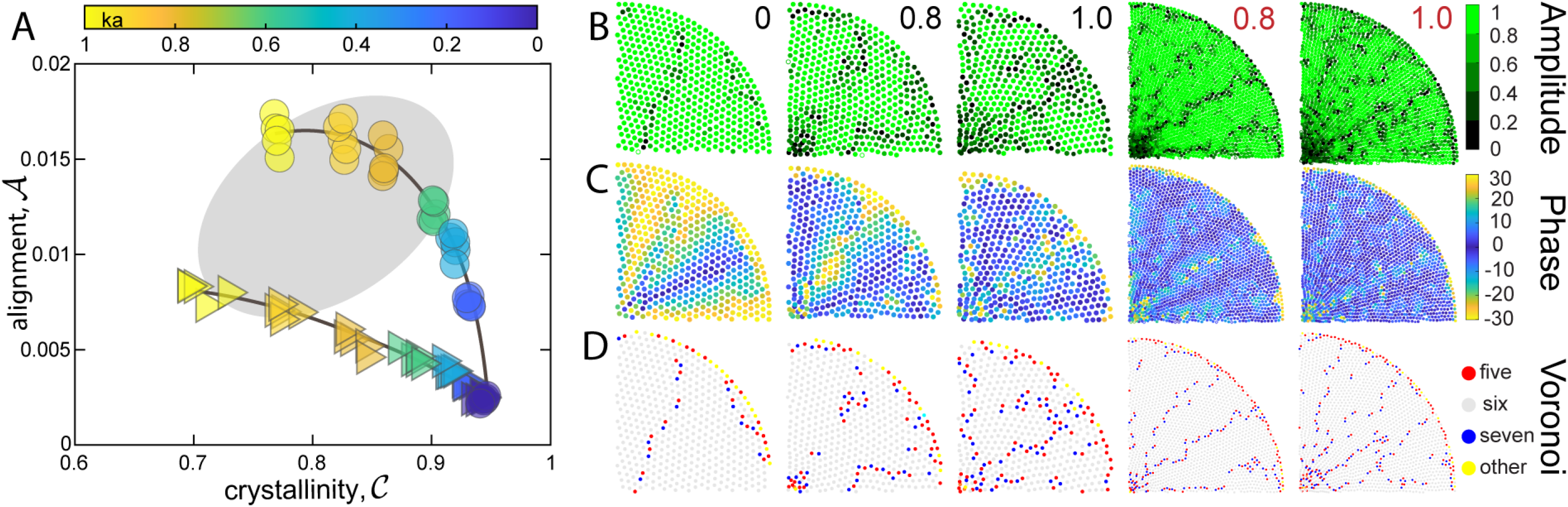
Simulations

**FIG. S8:**
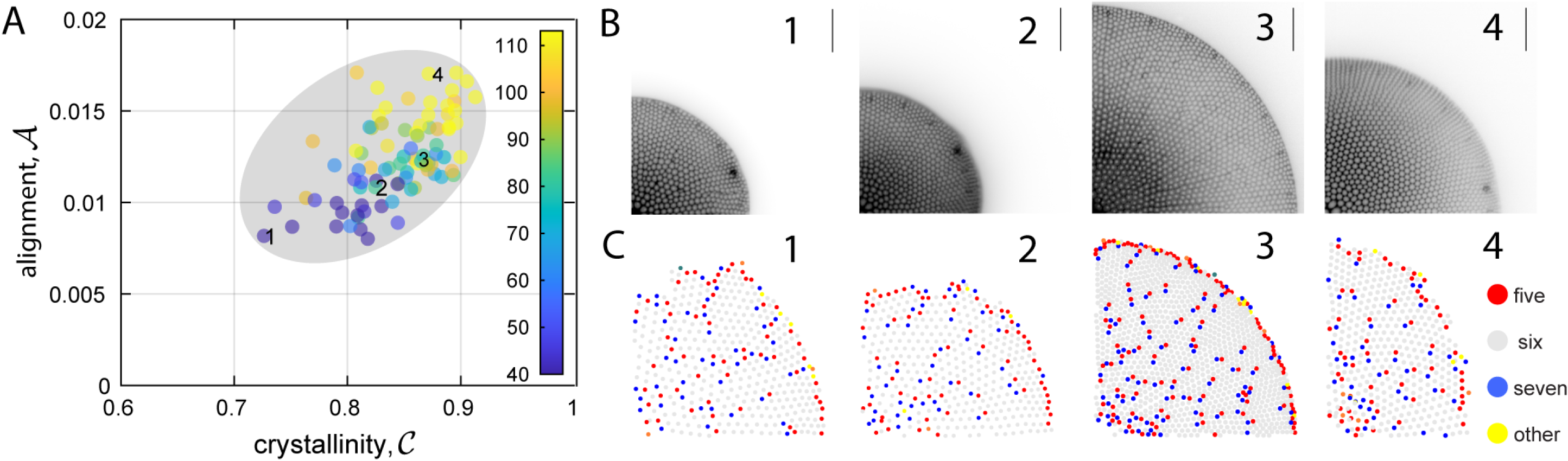
CG

